# An ancient antimicrobial protein co-opted by a fungal plant pathogen for *in planta* mycobiome manipulation

**DOI:** 10.1101/2021.06.10.447847

**Authors:** Nick C. Snelders, Gabriella C. Petti, Grardy C. M. van den Berg, Michael F. Seidl, Bart P.H.J. Thomma

**Affiliations:** Laboratory of Phytopathology, Wageningen University & Research, Wageningen, The Netherlands; Cluster of Excellence on Plant Sciences (CEPLAS), Institute for Plant Sciences, University of Cologne, Cologne, Germany; Theoretical Biology & Bioinformatics Group, Department of Biology, Utrecht University, Utrecht, The Netherlands

## Abstract

Microbes typically secrete a plethora of molecules to promote niche colonization. Soil-dwelling microbes are well-known producers of antimicrobials that are exploited to outcompete microbial co-inhabitants. Also plant pathogenic microbes secrete a diversity of molecules into their environment for niche establishment. Upon plant colonization, microbial pathogens secrete so-called effector proteins that promote disease development. While such effectors are typically considered to exclusively act through direct host manipulation, we recently reported that the soil-borne fungal xylem-colonizing vascular wilt pathogen *Verticillium dahliae* exploits effector proteins with antibacterial properties to promote host colonization through the manipulation of beneficial host microbiota. Since fungal evolution preceded land plant evolution, we now speculate that a subset of the pathogen effectors involved in host microbiota manipulation evolved from ancient antimicrobial proteins of terrestrial fungal ancestors that served in microbial competition prior to the evolution of plant pathogenicity. Here, we show that *V. dahliae* has co-opted an ancient antimicrobial protein as effector, named VdAMP3, for mycobiome manipulation *in planta*. We show that VdAMP3 is specifically expressed to ward off fungal niche competitors during resting structure formation in senescing mesophyll tissues. Our findings indicate that effector-mediated microbiome manipulation by plant pathogenic microbes extends beyond bacteria and also concerns eukaryotic members of the plant microbiome. Finally, we demonstrate that fungal pathogens can exploit plant microbiome-manipulating effectors in a life-stage specific manner, and that a subset of these effectors has evolved from ancient antimicrobial proteins of fungal ancestors that likely originally functioned in manipulation of terrestrial biota.

**SIGNIFICANCE STATEMENT:** Microbes secrete a diversity of molecules into their environment to mediate niche colonization. During host ingress, plant pathogenic microbes secrete effector proteins that facilitate disease development, many of which deregulate host immune responses. We recently demonstrated that plant pathogens additionally exploit effectors with antibacterial activities to manipulate beneficial plant microbiota to promote host colonization. Here, we show that the fungal pathogen *Verticillium dahliae* has co-opted an ancient antimicrobial protein, that likely served in microbial competition in terrestrial environments before land plants existed, as effector for the manipulation of fungal competitors during host colonization. Thus, we demonstrate that pathogen effector repertoires comprise antifungal proteins, and speculate such effectors could be exploited for the development of novel antimycotics.

## INTRODUCTION

Microbes are found in a wide diversity of niches on our planet. To facilitate establishment within microbial communities, microbes secrete a multitude of molecules to manipulate each other. Many of these molecules exert antimicrobial activities and are exploited to directly suppress microbial co-inhabitants in order to outcompete them for the limitedly available nutrients and space of a niche. Microbially-secreted antimicrobials encompass diverse molecules including peptides (AMPs) and lytic enzymes, but also non-proteinaceous molecules such as secondary metabolites. Soils are among the most biologically diverse and microbially competitive environments on earth. Microbial proliferation in the soil environment is generally limited by the availability of organic carbon (1), for which soil microbes continuously compete. Consequently, numerous saprophytic soil-dwelling microbes secrete potent antimicrobials that promote niche protection or colonization. Notably, these microbes are the primary source of our clinically used antibiotics (2, 3).

Like free-living microbes, also microbial plant pathogens secrete a multitude of molecules into their environment to mediate niche colonization (4, 5). The study of molecules secreted by microbial plant pathogens has been largely confined to the context of binary interactions between pathogens and hosts. To establish disease, plant pathogenic microbes secrete a plethora of so-called effectors, molecules of various kinds that promote host colonization and that are typically thought to mainly deregulate host immune responses (4, 6, 7). Upon host colonization, plant pathogens encounter a plethora of plant-associated microbes that collectively form the plant microbiota, which represents a key factor for plant health. Beneficial plant-associated microbes are found in and on all organs of the plant and help to mitigate (a)biotic stresses (8–13). Plants shape their microbiota and specifically attract beneficial microbes to suppress pathogens (14–16). Hence, the plant microbiome can be considered an inherent, exogenous layer that complements the plant’s endogenous innate immune system. We previously hypothesized that plant pathogens not only utilize effectors to target components of host immunity as well as other aspects of host physiology to support host colonization, but also to target the host microbiota in order to establish niche colonization (4, 5). We recently provided experimental evidence for this hypothesis by showing that the ubiquitously expressed effector VdAve1 that is secreted by the soil-borne fungal plant pathogen *Verticillium dahliae* acts as a bactericidal protein that promotes host colonization through the selective manipulation of host microbiomes by suppressing microbial antagonists (17, 18). Additionally, we demonstrated that VdAve1 and a further antibacterial effector named VdAMP2 are exploited by *V. dahliae* for microbial competition in soil and promote virulence of *V. dahliae* in an indirect manner (18). Collectively, these observations demonstrate that *V. dahliae* dedicates part of its effector catalog towards microbiota manipulation. Likely, the V. dahliae genome encodes further effectors that act in microbiome manipulation.

Evidently, bacterial and fungal evolution on land preceded land plant evolution. As a consequence, fungal pathogen effectors involved in the manipulation of (host-associated) microbial communities may have evolved from ancestors that served in microbial competition in terrestrial niches hundreds of millions of years ago prior to land plant evolution. However, evidence for this hypothesis is presently lacking.

*V. dahliae* is an asexual xylem-dwelling fungus that causes vascular wilt disease on hundreds of plant species (19). The fungus survives in the soil in the form of multicellular melanized resting structures, called microsclerotia, that offer protection against (a)biotic stresses and can persist in the soil for many years (20). Microsclerotia represent the major inoculum source of *V. dahliae* in nature and their germination is triggered by carbon- and nitrogen-rich exudates from plant roots (21). Following microsclerotia germination, fungal hyphae grow through the soil and rhizosphere towards the roots of host plants. Next, *V. dahliae* colonizes the root cortex and crosses the endodermis, from which it invades xylem vessels. Once the fungus enters those vessels it forms conidiospores that are transported with the water flow until they get trapped, for instance by vessel end walls. This triggers germination of the conidiospores, followed by penetration of cell walls, hyphal growth and renewed sporulation, leading to systematic colonization of the plant (22). Once tissue necrosis commences and plant senescence occurs, host immune responses fade and *V. dahliae* enters a saprophytic phase when it emerges from the xylem vessels to invade adjacent host tissues, which is accompanied by the production of microsclerotia. Upon littering and decomposition of plant tissues, these microsclerotia are released into the soil (23).

## RESULTS

To identify effectors potentially acting in microbiome manipulation, we recently queried the *V. dahliae* secretome for structural homologs of known antimicrobial proteins (AMPs), which led to the identification of ten candidates, including the functionally characterized VdAMP2 (18). Among the remaining nine candidates we now identified a small cysteine-rich protein of ~4.9 kDa, which we name VdAMP3. As a first step in the characterization of VdAMP3 we assessed its predicted structure. Interestingly, VdAMP3 is predicted to adopt a Cysteine-stabilized αβ (CSαβ) fold that is also found in defensin-like proteins (Fig. 1a)(24–26). CSαβ defensins represent a wide-spread and well-characterized family of antimicrobial proteins that are presumed to share a single ancient origin in the last common ancestor of animals, plants and fungi that produce these proteins today (24–27). It is important to note, however, that many typical small cysteine-rich pathogen effectors adopt AMP-like confirmations, and that tertiary structures of several AMP families strongly resemble each other (27, 28). Hence, structure prediction can easily lead to false-positive classifications as AMP or allocation to the wrong AMP family.

**Figure 1.**
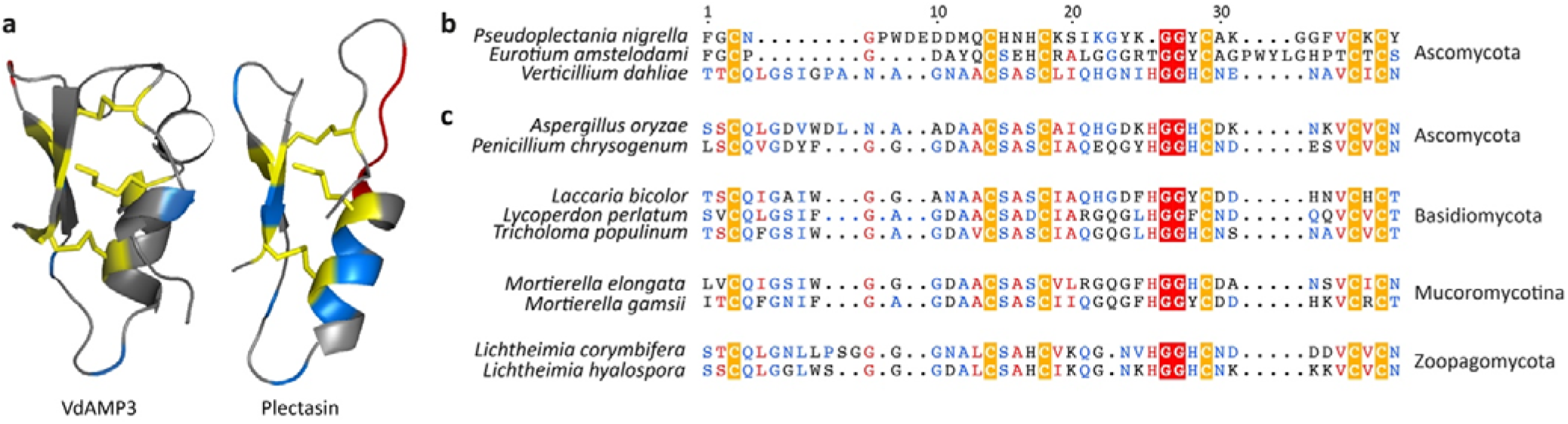
The *V. dahliae* effector VdAMP3 evolved from an ancient fungal protein. **(a)** VdAMP3 (left) is predicted to adopt a cysteine-stabilized αβ (CSαβ) defensin-like fold. The structure of the CSαβ defensin Plectasin (right) of the fungus *Pseudoplectania nigrella* is included as reference. The disulfide bonds stabilizing the antiparallel β-sheets and the α-helix are highlighted in yellow. Positively and negatively charged amino acid residues are highlighted in blue and red, respectively. **(b)** Protein sequence alignment with CSαβ defensins Plectasin and Eurocin (*Eurotium amstelodami*) supports the structure prediction of VdAMP3. **(c)** VdAMP3 homologs are widespread in the fungal kingdom. Protein sequence alignment of VdAMP3 with a subset of its homologs identified in higher (Ascomycota and Basidiomycota) and lower fungi (Mucoromycotina and Zoopagomycota). The alignment as shown in **(b-c)** displays the most conserved region of the CSαβ defensin protein family and was performed using HMMER and visualized with Espript3. The highly conserved cysteine and glycine residues that contribute to the CSαβ defensin structure are highlighted by yellow and red backgrounds, respectively. The homologs displayed in **(c)** were identified using blastP in the predicted proteomes of the respective fungi included in the JGI 1000 Fungal Genomes Project (32).

CSαβ defensins, or so-called cis-defensins, owe their structure to highly conserved *cis*-orientated disulfide bonds that establish an interaction between a double- or triple-stranded antiparallel β-sheet with an α-helix (25, 27). To validate the prediction of VdAMP3 as a member of this ancient antimicrobial protein family, we aligned its amino acid sequence with the antibacterial CSαβ defensins Plectasin and Eurocin, from the saprophytic Ascomycete species *Pseudoplectania nigrella* and *Eurotium amstelodami* (formerly *Aspergillus amstelodami*), respectively (29–31). Although the biological relevance of these defensins for the respective fungi remains unclear, their antibacterial activity and protein structure have been well characterized, which lead to their recognition as genuine CSαβ defensins (29–31). Although the overall identity between the three proteins was rather low (25-40%), protein sequence alignment revealed that VdAMP3 contains the six highly conserved cysteine residues that are considered crucial for the structure of CSαβ defensins (Fig. 1b)(27). To further substantiate the emerging picture that VdAMP3 belongs to this particular protein family, and that the detected similarities with Plectasin and Eurocin are not the result of convergent protein evolution, we queried the predicted proteomes of the fungi from the JGI 1000 Fungal Genomes Project (32) for homologs of VdAMP3 with higher sequence identity and included a subset of those in the protein alignment (Fig. 1c). Interestingly, besides homologs in Ascomycota and Basidiomycota, our sequence similarity search also revealed homologs in early-diverging fungi from the subphyla Mucoromycotina and Zoopagomycota (both formerly classified as Zygomycota (33)) (Fig. 1c). Importantly, this divergence is estimated to have taken place approximately 900 million years ago (34), indicating it preceded the evolution of the first land plants approximately 450 million years later (34–37). Consequently, this analysis indicates that *VdAMP3* evolved from an ancestral fungal gene hundreds of millions of years ago, before land plants existed.

As a first step to determine the role of VdAMP3 in *V. dahliae* infection biology, we assessed conditions for *VdAMP3* expression. Transcriptome analysis of diverse *V. dahliae* strains during colonization of a diversity of hosts did not reveal *in planta* expression of VdAMP3 thus far (17, 38–40). However, strong induction of this effector gene was reported during microsclerotia formation in a transcriptome analysis of *V. dahliae* strain XS11 grown *in vitro* (24). To validate this finding, we analyzed *in vitro* expression of *VdAMP3* in *V. dahliae* strain JR2. To this end, *V. dahliae* conidiospores were spread on nitrocellulose membranes placed on top of solid minimal medium and fungal material was harvested prior to microsclerotia formation, after 48 hours of incubation, and after the onset of microsclerotia formation, after 96 hours of incubation. Expression of *VdAMP3* was determined at both time points with real-time PCR alongside expression of the *Chr6g02430* gene that encodes a putative cytochrome P450 enzyme that acts as a marker for microsclerotia formation (24, 41). Consistent with the observations for *V. dahliae* strain XS11 (24), no *VdAMP3* expression was detected at 48 hours when also *Chr6g02430* was not expressed and no visual microsclerotia formation could be observed on the growth medium (Fig. 2a). However, induction of *VdAMP3* as well as *Chr6g02430* was observed after 96 hours of incubation, at which time point also the formation of microsclerotia on the growth medium became apparent (Fig. 2a). Collectively these data demonstrate that expression of *VdAMP3* coincides with microsclerotia formation *in vitro* also for *V. dahliae* strain JR2.

**Figure 2.**
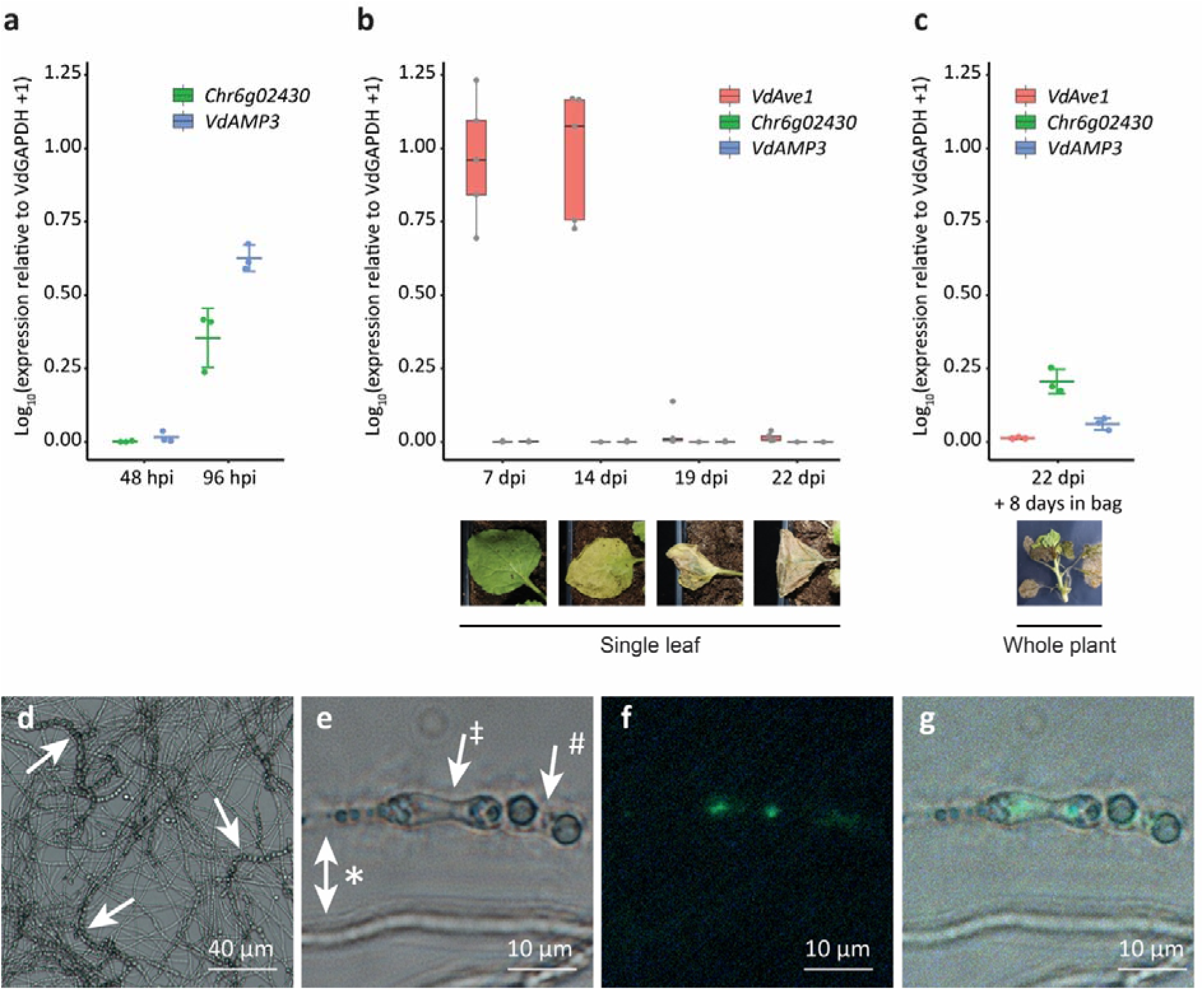
*VdAMP3* is specifically expressed in hyphal cells that develop into microsclerotia. **(a)** Expression of *VdAMP3* and the marker gene for microsclerotia development *Chr6g02430*, relative to the household gene *VdGAPDH* at 48 and 96 hours of *in vitro* cultivation (N=3). **(b)** Expression of *VdAve1*, *VdAMP3* and *Chr6g02430* in *N. benthamiana* leaves from 7 to 22 days post inoculation (dpi) (N=5). **(c)** Expression of *VdAve1*, *VdAMP3* and *Chr6g02430* in tissue of *N. benthamiana* plants harvested at 22 days post inoculation after 8 days of incubation in sealed plastic bags (N=3). **(d)** Microsclerotia formation of a *pVdAMP3::eGFP* reporter mutant as detected after 7 days of cultivation in Czapek Dox medium. Typical chains of microsclerotia (42, 43) are indicated by arrows. **(e)** Bright field image of various *V. dahliae* cell types after 7 days of cultivation in Czapek Dox, including hyphae (*), swollen hyphal cells developing into microsclerotia (‡) and mature microsclerotia cells (#). **(f)** GFP signal for the image as shown in **(e)**, indicative for activity of the *VdAMP3* promoter, is exclusively detected in the swollen hyphal cells developing into microsclerotia. **(g)** Overlay of **(e)** and **(f)**.

Although previous transcriptome analyses failed to detect *in planta* expression of *VdAMP3*, we realized that these analyses were predominantly performed for infection stages when the fungus is still confined to the xylem vessels and microsclerotia formation had not yet been initiated. Accordingly, *in planta* expression of **VdAMP3** may have been missed. Thus, we inoculated *Nicotiana benthamiana* with *V. dahliae* and determined expression of *VdAMP3* in leaves and petioles sampled at different time points and displaying different disease phenotypes, ranging from asymptomatic at seven days post inoculation (dpi) to complete necrosis at 22 dpi. As expected, a strong induction of the previously characterized *VdAve1* effector gene was detected at seven and 14 dpi (Fig. 2b) (17, 18). In contrast, however, no expression of *VdAMP3* was recorded, even at the latest time point when the leaf tissue had become completely necrotic (Fig. 2b). Importantly, also no *Chr6g02430* expression was detected at any of these time points (Fig. 2b), suggesting that microsclerotia formation had not yet started in these tissues. Indeed, visual inspection of the necrotic plant tissue collected at 22 dpi did not reveal microsclerotia presence. To induce microsclerotia formation, *V. dahliae*-inoculated *N. benthamiana* plants harvested at 22 dpi were sealed in plastic bags and incubated in the dark to increase the relative humidity and mimic conditions that occur during tissue decomposition in the soil. Interestingly, after eight days of incubation the first microsclerotia could be observed, and induction of *VdAMP3* as well as *Chr6g02430* was detected (Fig. 2c). Collectively, these findings suggest that *in planta* expression of *VdAMP3* coincides with microsclerotia formation, similar to our observations *in vitro*. Moreover, our data suggest that *VdAMP3* expression primarily depends on a developmental stage of *V. dahliae* rather than on host factors such as tissue necrosis.

To determine more precisely where *VdAMP3* is expressed, and to improve our understanding of how *V. dahliae* may benefit from effector expression during microsclerotia formation, we generated a *V. dahliae* reporter strain expressing eGFP under control of the *VdAMP3* promoter. Intriguingly, microscopic analysis of the reporter strain during microsclerotia formation stages *in vitro* (Fig. 2d), revealed that *VdAMP3* is expressed by swollen hyphal cells that act as primordia that subsequently develop into microsclerotia, but not by the adjacent hyphal cells or recently developed microsclerotia cells (Fig. 2e-g). This highly specific expression of *VdAMP3* suggests that the effector protein may facilitate the formation of microsclerotia in decaying host tissue. Given its presumed antimicrobial activity, *VdAMP3* may be involved in antagonistic activity against opportunistic decay organisms in this microbially competitive niche.

To determine if *VdAMP3* indeed exerts antimicrobial activity, we incubated a randomly selected panel of bacterial isolates with the effector protein and monitored their growth *in vitro*. VdAMP3 concentrations as high as 20 μM resulted in no or only marginal bacterial growth inhibition (Supplementary Fig. 1). A similar assay with fungal isolates showed that incubation with 5 μM of *VdAMP3* already markedly affected growth of the filamentous fungi *Alternaria brassicicola* and *Cladosporium cucumerinum* and the yeasts *Pichia pastoris* and *Saccharomyces cerevisiae* (Figure 3a). This finding suggests that *VdAMP3* displays more potent activity against fungi than against bacteria.

**Figure 3.**
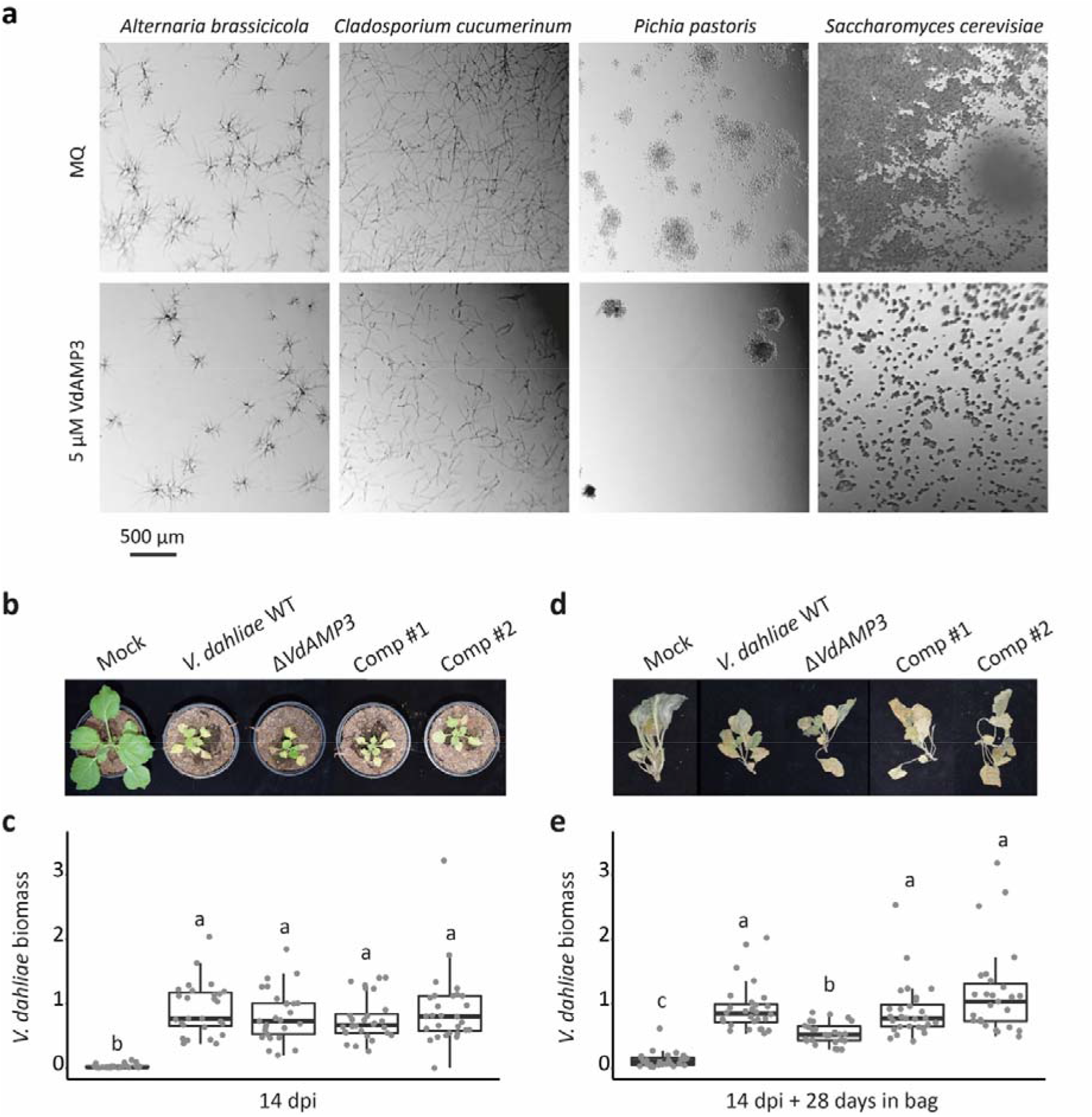
VdAMP3 is an antifungal protein that contributes to *V. dahliae* biomass accumulation in the decaying host phyllosphere. **(a)** Microscopic pictures of fungal isolates grown in 0.05× potato dextrose broth supplemented with 5 μM VdAMP3 or ultrapure water (MQ). VdAMP3 impairs growth of *Alternaria brassicicola, Cladosporium cucumerinum, Pichia pastoris* and *Saccharomyces cerevisiae*. Pictures were taken after 24 (*A. brassicicola, C. cucumerinum* and *S. cerevisiae*) or 64 (*P. pastoris*) hours of incubation. **(b)** VdAMP3 does not contribute to establishment of Verticillium wilt disease in *N. benthamiana*. Photos display representative phenotypes of *N. benthamiana* plants infected by wild-type *V. dahliae* (WT), the VdAMP3 deletion (Δ*VdAMP3*) and two complementation (Comp) mutants 14 days post inoculation. **(c)** Relative *V. dahliae* biomass in above-ground *N. benthamiana* tissues determined with real-time PCR. Different letter labels represent significant differences (one-way ANOVA and Tukey’s post-hoc test; p<0.05; N≥27 **(d)** Representative phenotypes of *N. benthamiana* plants as shown in **(b)** after 28 days of incubation in plastic bags. **(e)** Relative *V. dahliae* biomass in *N. benthamiana* tissues as displayed in **(d)**. Letters represent significant differences (one-way ANOVA and Tukey’s post-hoc test; p<0.05; N≥27).

To study the importance of the antifungal activity of VdAMP3 during and after host colonization, a *VdAMP3* deletion mutant was generated as well as complementation strains (Supplementary Fig 2). Importantly, targeted deletion of *VdAMP3* did not affect growth nor microsclerotia formation *in vitro* (Supplementary Fig. 3a,b). To determine if VdAMP3 contributes to Verticillium wilt disease development, *N. benthamiana* plants were inoculated with wild-type *V. dahliae* and the *VdAMP3* deletion mutant. In line with our inability to detect expression during early infection stages, disease phenotypes and *V. dahliae* biomass quantification using real-time PCR did not reveal a contribution of VdAMP3 to host colonization up to two weeks after inoculation (Fig. 3b,c). To test if VdAMP3 contributes to *V. dahliae* niche establishment following systemic host colonization, we harvested the *N. benthamiana* plants and sealed them in plastic bags to induce microsclerotia formation. Interestingly, following four weeks of incubation, *V. dahliae* biomass quantification in *N. benthamiana* plants inoculated with the various genotypes using real-time PCR revealed a significant reduction in biomass of the *VdAMP3* deletion mutant when compared with wild-type *V. dahliae* and complementation mutants (Fig 3d,e).

To investigate if the effects of VdAMP3 are limited to *N. benthamiana*, or whether those also extend to other hosts, we inoculated *Arabidopsis thaliana* plants with wild-type *V. dahliae* and the *VdAMP3* deletion mutant. Consistent with our observations for *N. benthamiana*, deletion of *VdAMP3* did not affect establishment of Verticillium wilt in *A. thaliana* (Supplementary Fig. 4a,b). However, *V. dahliae* biomass quantification in above-ground *A. thaliana* tissues at three weeks post inoculation revealed reduced accumulation of *V. dahliae* in the absence of VdAMP3 (Supplementary Fig. 4c). Thus, the effects of VdAMP3 are not restricted to a single host.

**Figure 4.**
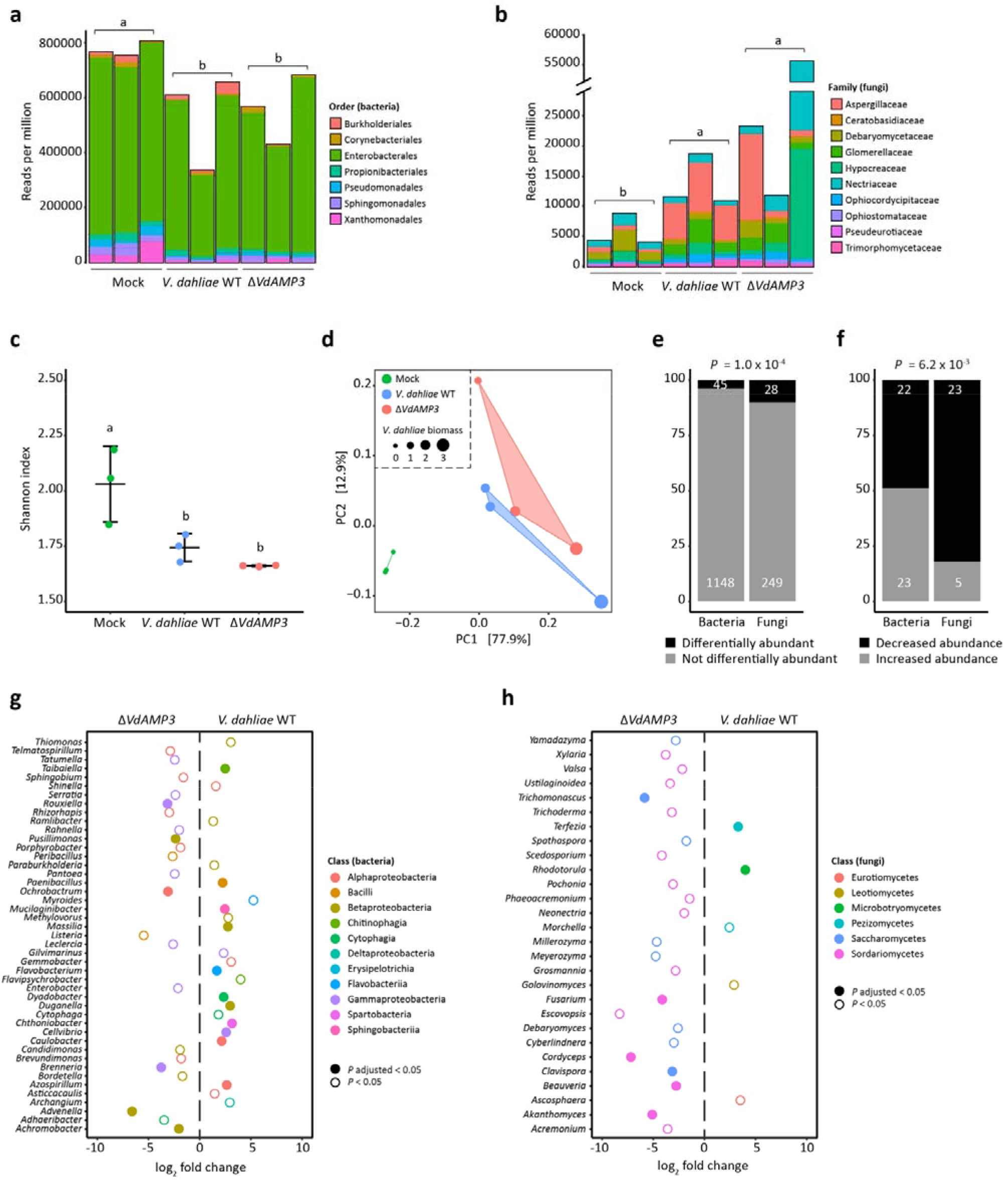
VdAMP3 manipulates the mycobiome of the decaying *N. benthamiana* phyllosphere. **(a-b)** *V. dahliae*-induced decay of the *N. benthamiana* phyllosphere is associated with a decreased bacterial, and increased fungal, abundance. Relative abundance of bacteria **(a)** and fungi **(b)**, excluding *V. dahliae*, in the phyllosphere of decaying *N. benthamiana* plants colonized by wild-type *V. dahliae* or the *VdAMP3* deletion mutant (14 days post inoculation and after 28 days of incubation in plastic bags), and in the phyllosphere of fresh *N. benthamiana* plants (mock). Letters represent significant differences in total bacterial/fungal abundance between the three treatments (one-way ANOVA and Tukey’s post hoc test; *P*⍰<⍰0.05; N=3). **(c)** *V. dahliae*-induced decay of *N. benthamiana* plants impacts alpha diversity of the phyllosphere. The plot displays the average Shannon index ± SD, letters represent significant differences (one-way ANOVA and Tukey’s post hoc test; *P*⍰<⍰0.05; N=3). **(d)** Principal coordinate analysis based on Bray-Curtis dissimilarities (beta diversity) reveals separation of the microbiomes based on the three different treatments. **(e)** Differential abundance analysis of microbial genera between the microbiomes colonized by *V. dahliae* WT and the *VdAMP3* deletion mutant indicates that secretion of *VdAMP3* significantly impacts a larger proportion of the fungi than of the bacteria (two-tailed Fisher’s exact test). **(f)** Of the differentially abundant microbial genera, significantly more fungi display a decreased abundance in the presence of *VdAMP3* when compared with the bacteria (two-tailed Fisher’s exact test). **(g-h)** Overview of the differentially abundant bacterial **(g)** and fungal **(h)** genera. The plots display increased (positive log_2_ fold change) or decreased (negative log_2_ fold change) abundance in the presence of *V. dahliae* WT when compared with the *VdAMP3* deletion mutant (Wald test, *P* adjusted < 0.05 and *P* < 0.05, N=3). Differentially abundant fungal genera from the Saccharomycetes or Sordariomycetes are consistently suppressed in the presence of VdAMP3.

As *in vitro* antimicrobial activity assays pointed towards fungi as the primary targets of *VdAMP3*, we speculated that *V. dahliae* exploits VdAMP3 to suppress fungal competitors in decomposing host tissues to safeguard the formation of its resisting structures. To characterize the microbiota associated with *N. benthamiana* decomposition and to determine the impact of VdAMP3 on these microbial communities, we characterized the phyllosphere microbiota of fresh mock-inoculated *N. benthamiana* plants, and decaying plants diseased by *V. dahliae* WT or the *VdAMP3* deletion mutant incubated in plastic bags, through shotgun metagenomic sequencing. Consistent with a primary role for fungi in the decomposition of dead plant material (44–48) we detected a significant increase of fungi and decrease of bacteria in the phyllosphere of the *N. benthamiana* plants diseased by the *V. dahliae* strains when compared with healthy mock-treated plants (Fig. 4a-b). These changes are accompanied by a reduced alpha diversity in the decaying phyllospheres (Fig. 4c). Additionally, principal coordinate analysis (PCoA) based on Bray-Curtis dissimilarities (beta diversity) uncovered clear separation of the microbiota of the healthy plants from those in decay (Fig. 4d). The PCoA also revealed a weaker, yet potentially relevant, separation of the microbiota colonized by *V. dahliae* WT and the *VdAMP3* deletion mutant, which suggests that secretion of *VdAMP3* manipulates microbiome compositions (Fig. 4d). Intriguingly, when we compared the abundances of the identified microbial genera between the microbiomes colonized by *V. dahliae* WT and the *VdAMP3* deletion mutant, we detected significantly more differentially abundant fungi (10.1%) than bacteria (3.8%) (Fig. 4e) (Supplementary Table 1-2). Interestingly, whereas the number of bacterial genera that display an increased or a decreased abundance in the presence of *VdAMP3* is more or less equal, the vast majority of the differentially abundant fungal genera (82.1%) are repressed in the presence of *VdAMP3* (Fig. 4f). Moreover, while no consistent suppression of bacterial genera from the same class could be detected, we exclusively identified suppression of the differentially abundant fungal genera from the Saccharomycetes or Sordariomycetes in the presence of *VdAMP3* (Fig. 4g-h). Thus, these observations indicate that *V. dahliae* VdAMP3 mainly acts as an antifungal effector protein that displays selective activity that predominantly impacts the mycobiome in the decaying host phyllosphere.

To further substantiate that the suppression of the Saccharomycetes and Sordariomycetes is a direct consequence of the *VdAMP3* activity, we incubated fungal species belonging to the suppressed genera with the effector to determine their sensitivity. In line with the previously observed sensitivity of the Saccharomycetes *. pastoris* and *S. cerevisiae*, also the Saccharomycete species *Cyberlindnera jadinii, Debaryomyces vanrijiae, Rhodotorula bogoriensis* and *Meyerozyma amylolytica* displayed markedly reduced growth in the presence of VdAMP3 (Fig. 5a, Supplementary Fig. 5). Similarly, growth of the Sordariomycetes *Cordyceps militaris* and *Trichoderma viride* was inhibited by the effector (Supplementary Fig. 5). Hence, these findings support the observed suppression of the Saccharomycetes and Sordariomycetes in the *N. benthamiana* phyllosphere mycobiome as a direct consequence of VdAMP3 activity.

**Figure 5.**
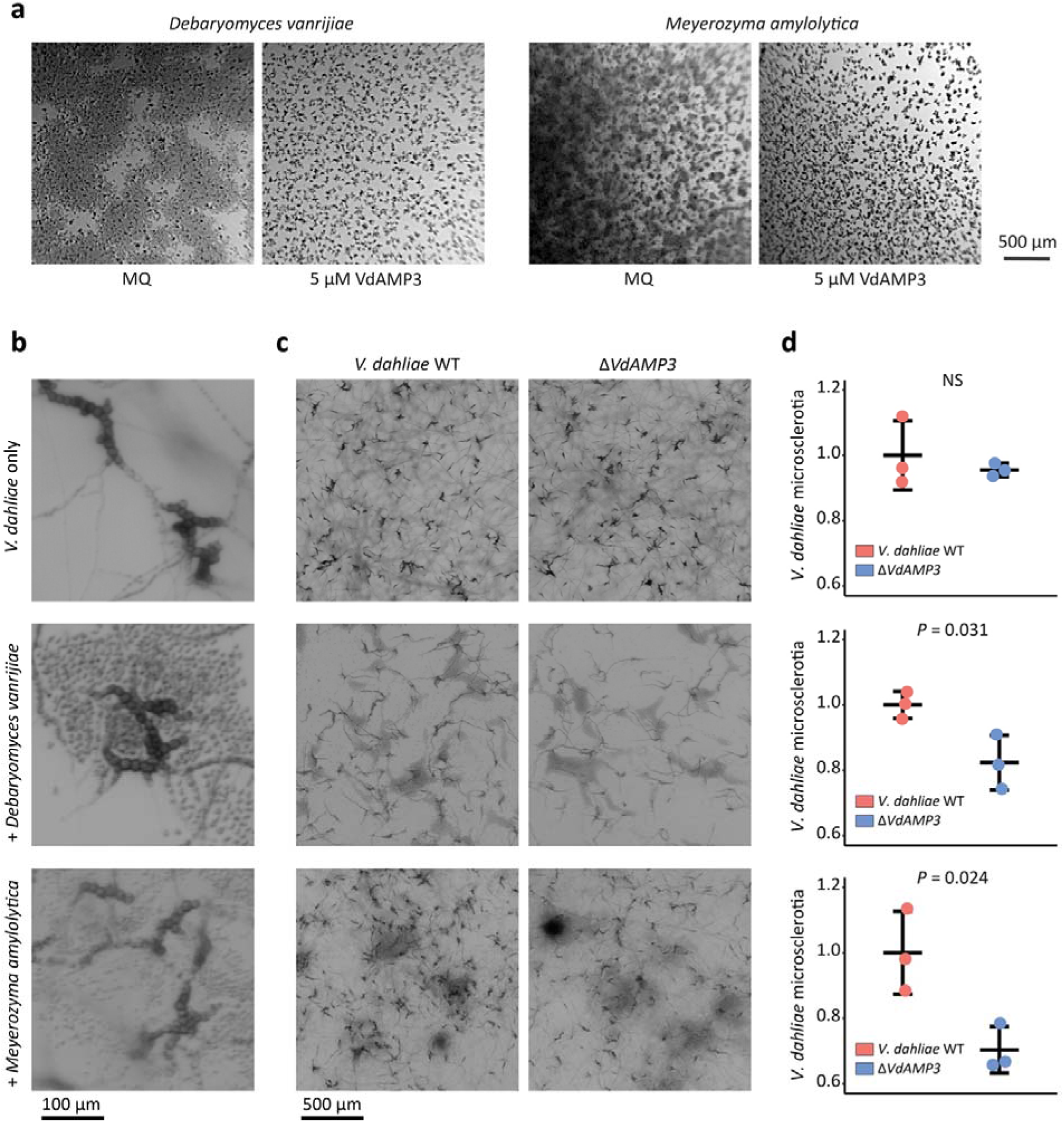
VdAMP3 contributes to *V. dahliae* microsclerotia formation in the presence of fungal niche competitors. **(a)** *Debaryomyces vanrijae* and *Meyerozyma amylolytica* are inhibited by VdAMP3. Microscopic pictures of the fungal species grown in 0.05x potato dextrose broth supplemented with 5 μM VdAMP3 or ultrapure water (MQ). Pictures were taken after 10 (*D. vanrijae*) or 24 (*Meyerozyma amylolytica*) hours of cultivation. **(b)** Close-up of *V. dahliae* microsclerotia formed after seven days of cultivation in the presence of *D. vanrijae* or *M. amylolytica*. **(c)** VdAMP3 contributes to *V. dahliae* microsclerotia formation in the presence of the other fungal species. Representative microscopic pictures displaying the co-culture of *V. dahliae* with *D. vanrijae* or *M. amylolytica*. Pictures were taken after seven days of co-cultivation. **(d)** Relative number of microsclerotia formed by *V. dahliae* WT and the *VdAMP3* deletion mutant in the presence of *D. vanrijae* or *M. amylolytica* as determined using ImageJ (unpaired two-sided student’s t-test; N=3).

The cell type-specific expression of VdAMP3, combined with its role in mycobiome manipulation, strongly suggests that VdAMP3 is exploited to ward off fungal niche competitors *in planta* to safeguard the formation of *V. dahliae* microsclerotia. To test if VdAMP3 indeed is essential for *V. dahliae* microsclerotia formation in the presence of other fungi, we co-cultivated *V. dahliae* WT and the *VdAMP3* deletion mutant with *D. vanrijiae* and *M. amylolytica*. Once microsclerotia formation by *V. dahliae* WT became apparent (Fig. 5b), we quantified the number of resting structures that were formed when compared with the *VdAMP3* deletion mutant. As anticipated, we detected a significant reduction of microsclerotia formed by the *VdAMP3* deletion mutant in the presence of both fungal species, confirming that *V. dahliae* relies on the antifungal activity of *VdAMP3* to form microsclerotia in the presence of particular fungal niche competitors (Fig. 5c,d).

## DISCUSSION

Microbes secrete a plethora of molecules to promote niche colonization (4). Free-living microbes are well known producers of antimicrobials that are secreted to outcompete microbial co-inhabitants to establish themselves in a microbial community. Microbial plant pathogens secrete a diversity of so-called effector molecules during host ingress, many of which are small cysteine-rich proteins that deregulate host immune responses to promote colonization (4, 6, 7). While investigating the vascular wilt fungus *V. dahliae*, we recently demonstrated that plant pathogens not only exploit effector proteins to promote disease establishment through direct host manipulation, but also through the manipulation of plant microbiota by means of antibacterial activities (18). Considering that the advent of fungi on earth preceded land plant evolution, we speculated that a subset of the pathogen effectors involved in host microbiota manipulation may have evolved from antimicrobial proteins that originally functioned in microbial competition in terrestrial niches before the first land plants appeared and plant pathogenicity evolved. Here, we demonstrated that the soil-borne fungal plant pathogen *V. dahliae* has co-opted an ancient antimicrobial protein as effector for mycobiome manipulation *in planta* to safeguard the formation of its resting structures. Thus, our findings indicate that plant pathogenicity in fungi is not exclusively associated with the evolution of novel effectors that manipulate plants or their associated microbial communities, but also with the co-option of previously evolved secreted proteins that initially served alternative lifestyles, such as saprotrophism, as effectors to promote host colonization. Moreover, our findings indicate that effector-mediated manipulation of plant microbiota by microbial plant pathogens is not confined to bacterial targets, but extends to eukaryotic microbes.

Functional characterization of VdAMP3 unveiled that the effector evolved to play a life stage-specific role in microbiome-manipulation during microsclerotia formation by *V. dahliae*. Recently, we described the characterization of the first microbiome-manipulating effectors secreted by *V. dahliae*; VdAve1 and VdAMP2 (18). VdAve1 is a ubiquitously expressed bactericidal effector that promotes *V. dahliae* host colonization through the selective manipulation of host microbiota in the roots as well as in the xylem by suppressing microbial antagonists. Moreover, VdAve1 is also expressed in the soil biome where it similarly contributes to niche colonization. Intriguingly, VdAMP2 is exclusively expressed in soil, and like VdAve1 exerts an antibacterial activity that contributes to niche establishment. Interestingly, VdAMP2 and VdAve1 display divergent activity spectra, and therefore likely complement each other for optimal soil colonization. In decaying host tissue neither VdAve1 nor VdAMP2 is expressed, yet VdAMP3 expression occurs. Collectively, our findings for VdAve1, VdAMP2 and VdAMP3 demonstrate that *V. dahliae* dedicates a substantial part of its catalog of effector proteins towards microbiome manipulation, and that each of these effectors act in a life stage-specific manner.

The life stage-specific exploitation of the in planta secreted antimicrobial effectors VdAve1 and VdAMP3 is well reflected by their antimicrobial activities and by the microbiota of the niches where they act. Contrary to previous *V. dahliae* transcriptome analyses, that repeatedly identified *VdAve1* as one of the most highly expressed effector genes *in planta* (17, 38–40), we detected a repression of the effector gene in decomposing *N. benthamiana* tissues (Fig. 1b,c). Characterization of the antimicrobial activity exerted by VdAve1 previously uncovered that the protein exclusively affects bacteria and does not impact fungi (18). Thanks to their ability to produce a wide diversity of hydrolytic enzymes, fungi are the primary decomposers of plant debris on earth (44). The phyllosphere of plants comprises a diversity of fungi (49–51). Importantly, upon plant senescence, these fungi are provided the first access to decaying material on which they can act opportunistically once host immune responses have faded. Accordingly, we detected an increased abundance of fungi in the phyllosphere of the decomposing *N. benthamiana* plants diseased by *V. dahliae* when compared with healthy plants (Fig. 4b). The observed repression of *VdAve1* and the subsequent induction of *VdAMP3* in a niche where *V. dahliae* encounters more fungal competition, underscores the notion that *V. dahliae* tailors the expression of its microbiome-manipulating effectors according to the various microbiota that it encounters during the different life stages. Along these lines it is tempting to speculate that during saprotrophism in soil *V. dahliae* exploits antimicrobial effector proteins to ward off other eukaryotic competitors including soil-dwelling parasites such as fungivorous nematodes or protists. However, evidence for this hypothesis is presently lacking.

Antimicrobial resistance in bacteria and fungi is posing an increasing threat to human health. Possibly, microbiome-manipulating effectors represent a valuable source for the identification and development of novel antimicrobials that can be deployed to treat microbial infections. Arguably, our findings that microbiome-manipulating effectors secreted by plant pathogens also comprise antifungal proteins opens up opportunities for the identification and development of novel antimycotics. Most fungal pathogens of mammals are saprophytes that generally thrive in soil or decaying organic matter, but can opportunistically cause disease in immunocompromised patients (52–54). Azoles are an important class of antifungal agents that are used to treat fungal infections in humans. Unfortunately, agricultural practices involving massive spraying of azoles to control fungal plant pathogens, but also the extensive use of azoles in personal care products, ultraviolet stabilizers, and anti-corrosives in aircrafts, for instance, give rise to an enhanced evolution of azole resistance in opportunistic pathogens of mammals in the environment (52, 55). For instance, azole resistant *Aspergillus fumigatus* strains are ubiquitous in agricultural soils and in decomposing crop waste material where they thrive as saprophytes (56, 57). Thus, fungal pathogens of mammals, like *A. fumigatus*, comprise niche competitors of fungal plant pathogens. Hence, we speculate that, like *V. dahliae*, also other plant pathogenic fungi may carry potent antifungal proteins in their effector catalogues that aid in niche competition with these fungi. Possibly, the identification of such effectors could contribute to the development of novel antimycotics.

## MATERIALS AND METHODS

### Gene expression analyses

*In vitro* cultivation of *V. dahliae* strain JR2 for analysis of *VdAMP3* and *Chr6g02430* expression was performed as described previously (24). Additionally, for *in planta* expression analyses, total RNA was isolated from individual leaves or complete *N. benthamiana* plants harvested at different time points after *V. dahliae* root dip inoculation. To induce microsclerotia formation, *N. benthamiana* plants were harvested at 22 dpi and incubated in sealed plastic bags (volume = 500 mL) for 8 days, prior to RNA isolation. RNA isolations were performed using the Maxwell^®^ 16 LEV Plant RNA Kit (Promega, Madison, USA). Real-time PCR was performed as described previously using the primers listed in Supplementary Table 3 (17).

### Generation of *V. dahliae* mutants

The *VdAMP3* deletion and complementation mutants, as well as the eGFP expression mutant, were generated as described previously using the primers listed in Supplementary Table 3 (18). To generate the *VdAMP3* complementation construct, the *VdAMP3* coding sequence was amplified with flanking sequences (~0.9 kb upstream and ~0.8 kb downstream) and cloned into pCG (58). Finally, the construct was used for *Agrobacterium tumefaciens*-mediated transformation of *V. dahliae* as described previously (59). *in vitro* growth and microsclerotia production of the *VdAMP3* deletion mutant was tested and quantified as described previously (18).

### Microbial isolates

Bacterial strains *B. subtilis* AC95, *S. xylosus* M3, *P. corrugata* C26, *Streptomyces* sp. NE-P-8 and *Ralstonia* sp. M21 were obtained from our in-house endophyte culture collection. Bacterial strains *Novosphingobium* sp. (NCCB 100261) and *Sphingobacterium canadese* (NCCB100125) were obtained from the Westerdijk Fungal Biodiversity Institute (Utrecht, the Netherlands). Fungal strains *Saccharomyces cerevisiae* H15 and *Trichoderma viride* were obtained from our in-house culture collection. Fungal strains *Cyberlindnera jadinii* (DSM 70167), *Cordyceps militaris* (DSM 1153), *Debaryomyces vanrijiae* (DSM 70252), *Meyerozyma amylolytica* (DSM 27310) and *Rhodotorula bogoriensis* (DSM 70872) were obtained from the Leibniz Institute DSMZ.

### *In vitro* microbial growth assays

Bacterial isolates were grown on lysogeny broth agar (LBA) at 28 °C. Single colonies were selected and grown overnight in low salt LB (10 g/L tryptone, 5 g/L yeast extract and 0.5 g/L sodium chloride) at 28 °C while shaking at 200 rpm. Overnight cultures were resuspended to OD_600_=0.025 in fresh low salt LB supplemented with 20 μM VdAMP3 or ultrapure water (MQ). *In vitro* growth was quantified using a CLARIOstar plate reader (BMG Labtech) as described previously (18).

Fungal isolates were grown on potato dextrose agar (PDA) at 22 °C. For yeasts, single colonies were selected and grown overnight in 0.05x potato dextrose broth (PDB) at 28 °C while shaking at 200 rpm. Overnight cultures were resuspended to OD_600_=0.01 in fresh 0.05x potato dextrose broth supplemented with 5 μM VdAMP3 or ultrapure water (MQ). Alternatively, for filamentous fungi, spores were harvested from PDA and suspended in 0.05x potato dextrose broth supplemented with 5 μM VdAMP3 or ultrapure water (MQ) to a final concentration of 10^4^ spores/mL. Next, 200 μL of the fungal suspensions was aliquoted in clear 96-well flat-bottom polystyrene tissue culture plates. Plates were incubated at 28 °C and fungal growth was imaged using a SZX10 stereo microscope (Olympus) with EP50 camera (Olympus).

### Inoculation assays

Three-week-old *N. benthamiana* seedlings grown in the greenhouse at 21 °C/19 °C during 16h/8h day/night periods, respectively, with 70% relative humidity, were inoculated with *V. dahliae* through root-dip inoculation as described previously (60). After 14 days, above-ground parts of the *N. benthamiana* plants were harvested and stored at −20 °C. Alternatively, above-ground parts were collected and transferred to plastic bags (volume = 500 mL) and incubated for four weeks at room temperature. Next, all *N. benthamiana* samples were ground using mortar and pestle. Subsequent genomic DNA isolation and *V. dahliae* biomass quantification was performed as previously described using the primers listed in Supplementary Table 3 (61).

### Fluorescence microscopy

Conidiospores of the *pVdAMP3::eGFP* reporter strain were harvested from a PDA plate and diluted to a final concentration of 10^5^ conidiospores/mL in 0.1x Czapek Dox medium. The suspension was incubated for one week at room temperature to allow hyphae to grow and microsclerotia to form. Finally, eGFP accumulating in the fungal cells was detected using a Nikon ECLIPSE 90i microscope.

### Microbiome analysis

Inoculation and incubation of *N. benthamiana* plants was performed as described above. After four weeks of incubation in plastic bags at room temperature in the dark, the decaying *N. benthamiana* phyllosphere samples colonized by *V. dahliae* WT and the *VdAMP3* deletion mutant were collected. The phyllospheres of fresh three-week-old *N. benthamiana* plants were included as controls. All samples were flash-frozen in liquid nitrogen and ground using mortar and pestle, genomic DNA was isolated using the DNeasy PowerSoil Kit (Qiagen, Venlo, The Netherlands). Sequencing libraries were prepared using the TruSeq DNA Nano kit (Illumina, San Diego, CA) and paired-end 150 bp sequencing was performed on the Illumina NextSeq500 platform at the Utrecht Sequencing Facility (USEQ).

The sequencing data was processed as follows. Quality control of the reads, adapter trimming and removal of *N. benthamiana* reads was performed with the ATLAS metagenomic workflow using the default parameters of the configuration file (62). Reads of the different samples were combined and assembled using metaSPAdes (used k-mer sizes: 21, 33, 55) to obtain a single metagenome cross-assembly (63). Subsequently, the cross-assembled contigs were taxonomically classified using CAT and binned per genus (64). The reads of the individual samples were mapped to the binned contigs using BWA-MEM (65). Next, the mapping files were converted to bam-format using SAMtools (66) v1.10 and the number of reads mapped to the contigs of a single genus were converted to “reads per million” for the individual samples. The generated taxonomy table and abundance table were subsequently transformed into a phyloseq (67) object (v.1.30.0) in R (v.3.6.1) to facilitate analysis of the microbiomes. The alpha diversity (Shannon index) and beta diversity (Bray–Curtis dissimilarity) were determined as described previously (67, 68). The DESeq2 extension of phyloseq was used to identify differentially abundant microbial genera (69). To this end, a parametric model was applied to the data and a negative binomial Wald test was used to test for significant differential abundance.

### Fungal co-cultivation assays

Fungal isolates were grown on PDA at room temperature. For *D. vanrijiae* and *M. amylolytica* single colonies were selected and grown overnight in 0.05x PDB at 28°C while shaking at 200 rpm. The overnight cultures of *D. vanrijiae* and *M. amylolytica* were resuspended to OD_600_=0.001 and 0.0001 in fresh 0.05x PDB, respectively. Conidiospores of *V. dahliae* strain JR2 and the *VdAMP3* deletion mutant were harvested from PDA plates and diluted in ultrapure water (MQ) to a final concentration of 10^4^ conidiospores/mL. Next, 150 μL of the yeast suspensions were mixed with 150 μL of the *V. dahliae* condiospore suspensions in clear 24-well flat-bottom polystyrene tissue culture plates. Finally, after seven days of incubation at 22°C, fungal growth was imaged using a SZX10 stereo microscope (Olympus) with EP50 camera (Olympus). The number of microsclerotia formed by *V. dahliae* WT and the *VdAMP3* deletion mutant was quantified using ImageJ.

## Acknowledgements

B.P.H.J.T. is supported by the Research Council Earth and Life Sciences (ALW) of the Netherlands Organization of Scientific Research (NWO). B.P.H.J.T acknowledges funding by the Alexander von Humboldt Foundation in the framework of an Alexander von Humboldt Professorship endowed by the German Federal Ministry of Education and research is furthermore supported by the Deutsche Forschungsgemeinschaft (DFG, German Research Foundation) under Germany’s Excellence Strategy – EXC 2048/1 – Project ID: 390686111. We thank Utrecht Sequencing Facility, subsidized by the University Medical Center Utrecht, Hubrecht Institute, Utrecht University, and The Netherlands X-omics Initiative (NWO project 184.034.019), for providing sequencing service. The authors declare no competing interests exist.

## Author contributions

N.C.S. and B.P.H.J.T. conceived the project. N.C.S., G.C.P. and B.P.H.J.T. designed the experiments. N.C.S., G.C.P. and G.C.M.B. carried out the experiments. N.C.S., G.C.P., M.F.S. and B.P.H.J.T. analyzed the data. N.C.S. and B.P.H.J.T. wrote the manuscript. All authors read and approved the final manuscript.

## Data and materials availability

The metagenomics data have been deposited in the NCBI GenBank database under BioProject PRJNA728211.

