## Supplemental Data for "An ancient antimicrobial protein co-opted by a fungal plant pathogen for *in planta* mycobiome manipulation"

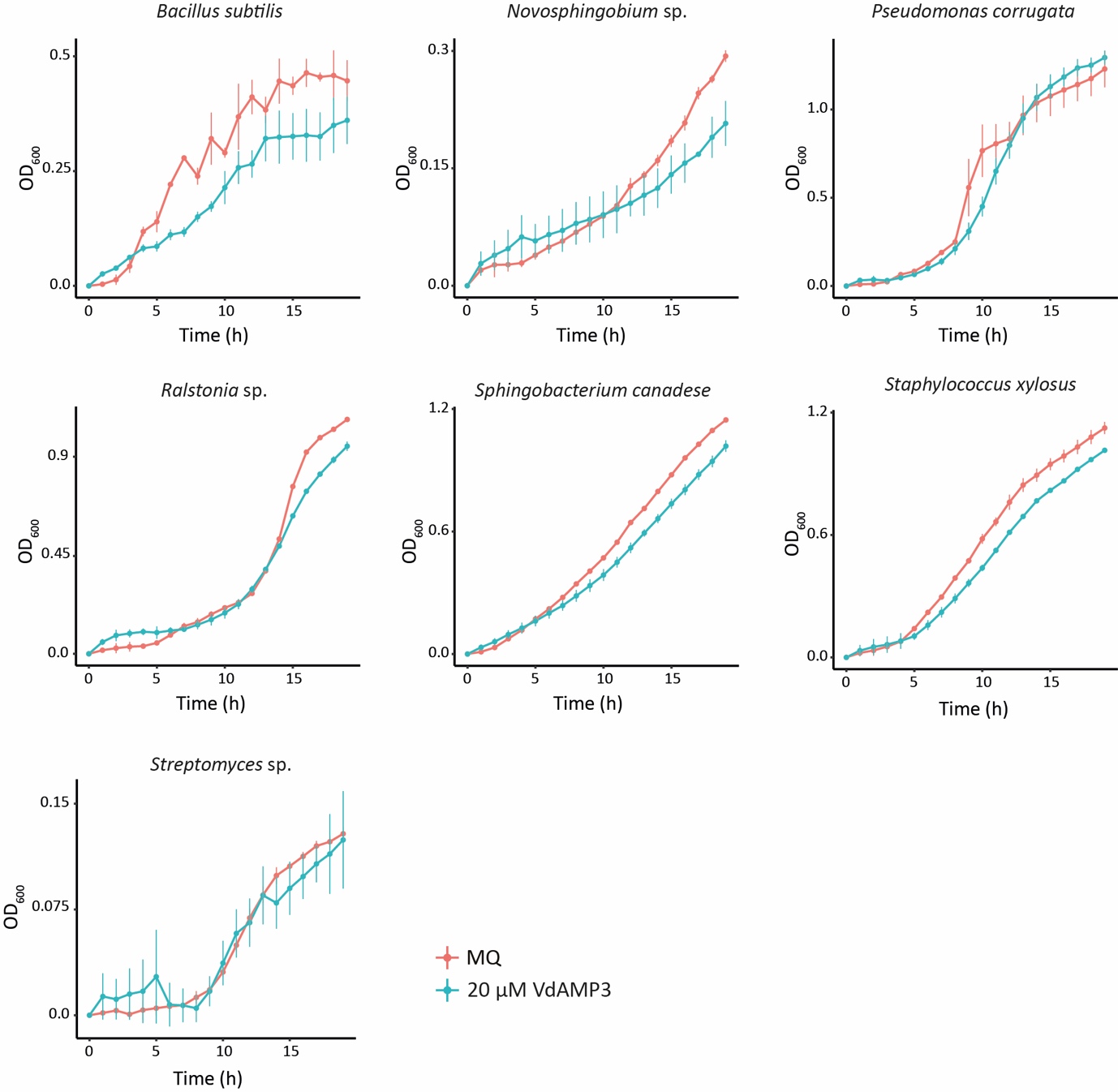

**Supplementary Figure 1. VdAMP3 does not markedly impact bacterial growth.** *In vitro* growth of plant-associated bacterial isolates in low salt LB is not, or only marginally, affected in the presence of 20 μM VdAMP3.

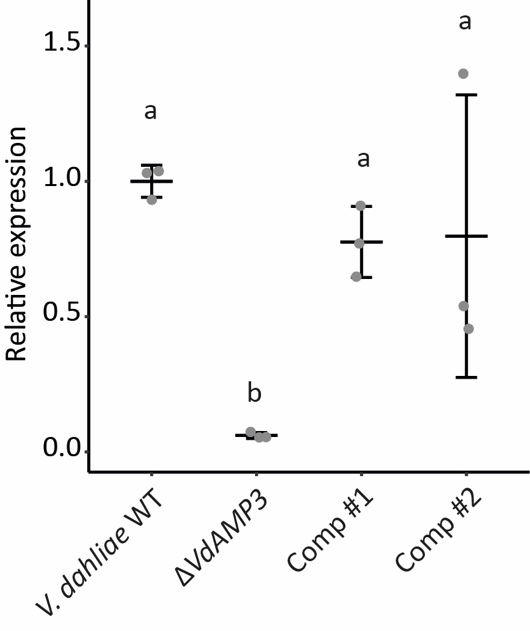

**Supplementary Figure 2. Expression of *VdAMP3* in *V. dahliae* mutants.** Expression of *VdAMP3* relative to *VdGAPDH* during microsclerotia formation by *V. dahliae* WT and the *VdAMP3* deletion and complementation mutants after seven days of cultivation in Czapek Dox medium. Letters represent statistically significant differences (one-way ANOVA and Tukey’s post-hoc test; p<0.05; N=3).

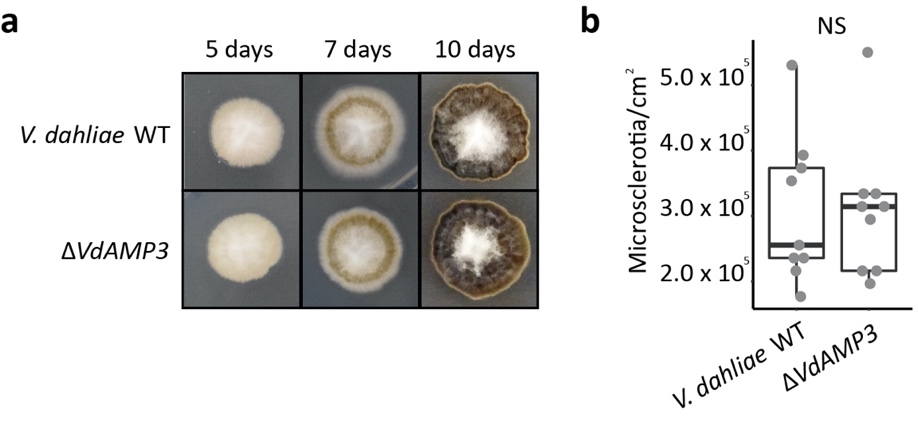

**Supplementary Figure 3. Deletion of *VdAMP3* does not affect *V. dahliae* microsclerotia formation *in vitro*. (a)** Morphology of wild-type *V. dahliae* and the *VdAMP3* deletion mutant following five, seven and ten days of *in vitro* growth on PDA. **(b)** Deletion of *VdAMP3* does not impair microsclerotia formation. After ten days, the colonies as shown in **(a)** were processed and the number of microsclerotia per cm^2­^ was determined using a haemocytometer. No significant difference in microsclerotia formation was observed (unpaired student t-test N=9).

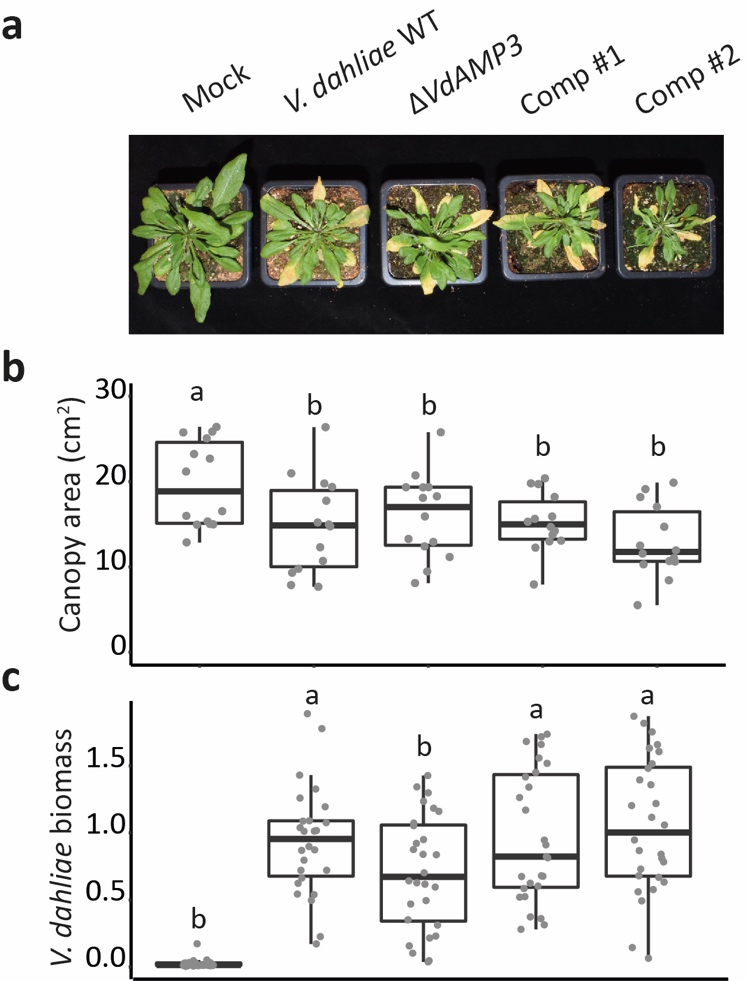

**Supplementary Figure 4. VdAMP3 contributes to *V. dahliae* biomass accumulation in *Arabidopsis thaliana* but does not influence development of disease symptoms. (a)** Deletion of *VdAMP3* does not affect establishment of Verticillium wilt disease in *A. thaliana*. Photos display representative phenotypes of *A. thaliana* plants 21 days post inoculation with *V. dahliae* WT and *VdAMP3* deletion and complementation mutants. **(b)** Canopy area of *A. thaliana* plants inoculated by the different *V. dahliae* genotypes. Different letter labels represent statistically significant differences when compared with *V. dahliae* WT (unpaired student’s t-test; N=14). **(c)** Relative *V. dahliae* biomass in above-ground *A. thaliana* tissues determined with real-time PCR. Different letter labels represent statistically significant differences when compared with *V. dahliae* WT (unpaired student’s t-test; N≥26).

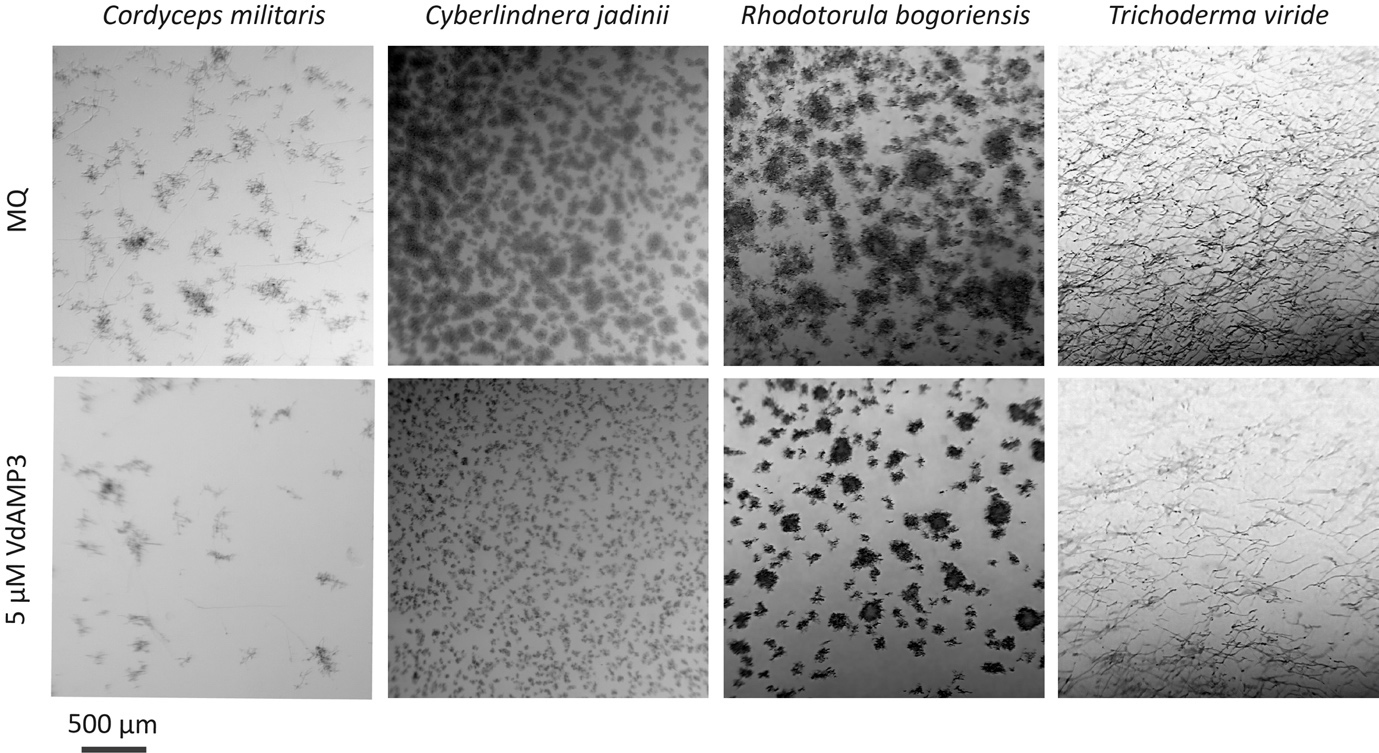

**Supplementary Figure 5. VdAMP3 affects Saccharomycetes and Sordariomycetes.** Microscopic pictures of fungal isolates grown in 0.05x potato dextrose broth supplemented with 5 µM VdAMP3 or ultrapure water (MQ). VdAMP3 impairs growth of *Cordyceps militaris*, *Cyberlindnera jadinii*, *Rhodotorula bogoriensis* and *Trichoderma viride*. Pictures were taken after 10 *(C. jadinii*), 24 (*R. bogoriensis*) or 30 (*C. militaris* and *T. viride*) hours of incubation.

**Supplementary Table 1: DESeq2 output for the differentially abundant bacterial genera in the decaying phyllosphere of *N. benthamiana* plants colonized by *V. dahliae* wild type compared to the *VdAMP3* deletion mutant**

| **Base mean** | **log2FC** | **lfcSE** | **stat** | **p value** | **p adjusted** | **Genus** |
| --- | --- | --- | --- | --- | --- | --- |
| 6.57E+03 | -6.60E+00 | 1.13E+00 | -5.82E+00 | 5.88E-09 | 2.26E-06 | *Advenella* |
| 1.43E+01 | -2.17E+01 | 3.91E+00 | -5.54E+00 | 2.99E-08 | 5.76E-06 | *Absicoccus* |
| 8.83E+01 | 2.65E+00 | 6.25E-01 | 4.23E+00 | 2.31E-05 | 2.97E-03 | *Azospirillum* |
| 3.87E+01 | 2.97E+00 | 7.72E-01 | 3.85E+00 | 1.18E-04 | 6.00E-03 | *Duganella* |
| 8.74E+01 | 3.16E+00 | 8.36E-01 | 3.78E+00 | 1.56E-04 | 6.00E-03 | *Chthoniobacter* |
| 9.34E+03 | 2.76E+00 | 7.30E-01 | 3.78E+00 | 1.59E-04 | 6.00E-03 | *Massilia* |
| 3.16E+02 | -3.09E+00 | 8.23E-01 | -3.76E+00 | 1.71E-04 | 6.00E-03 | *Ochrobactrum* |
| 5.66E+01 | -3.76E+00 | 1.08E+00 | -3.47E+00 | 5.13E-04 | 1.52E-02 | *Brenneria* |
| 2.74E+01 | 2.24E+00 | 6.71E-01 | 3.34E+00 | 8.37E-04 | 2.12E-02 | *Paenibacillus* |
| 5.08E+01 | -3.14E+00 | 9.45E-01 | -3.33E+00 | 8.82E-04 | 2.12E-02 | *Rouxiella* |
| 1.32E+03 | 2.15E+00 | 6.62E-01 | 3.25E+00 | 1.15E-03 | 2.61E-02 | *Caulobacter* |
| 5.62E+02 | 2.59E+00 | 8.32E-01 | 3.11E+00 | 1.87E-03 | 3.99E-02 | *Cellvibrio* |
| 9.76E+02 | 2.46E+00 | 7.94E-01 | 3.09E+00 | 1.97E-03 | 3.99E-02 | *Mucilaginibacter* |
| 1.85E+01 | -3.47E+00 | 1.13E+00 | -3.08E+00 | 2.10E-03 | 4.04E-02 | *Adhaeribacter* |
| 7.36E+02 | 1.68E+00 | 5.55E-01 | 3.02E+00 | 2.50E-03 | 4.58E-02 | *Flavobacterium* |
| 6.39E+01 | 2.49E+00 | 8.28E-01 | 3.01E+00 | 2.63E-03 | 4.59E-02 | *Taibaiella* |
| 2.72E+03 | 2.34E+00 | 7.89E-01 | 2.97E+00 | 2.98E-03 | 4.59E-02 | *Dyadobacter* |
| 5.20E+01 | -2.35E+00 | 7.98E-01 | -2.94E+00 | 3.27E-03 | 4.84E-02 | *Pusillimonas* |
| 2.76E+03 | 1.43E+00 | 5.02E-01 | 2.84E+00 | 4.46E-03 | 6.13E-02 | *Paraburkholderia* |
| 1.21E+01 | -2.97E+00 | 1.06E+00 | -2.79E+00 | 5.25E-03 | 6.60E-02 | *Rhizorhapis* |
| 2.26E+01 | -2.87E+00 | 1.08E+00 | -2.65E+00 | 7.97E-03 | 8.98E-02 | *Telmatospirillum* |
| 4.21E+00 | 5.25E+00 | 1.98E+00 | 2.65E+00 | 8.03E-03 | NA | *Myroides* |
| 5.77E+03 | -2.03E+00 | 7.67E-01 | -2.65E+00 | 8.15E-03 | 8.98E-02 | *Achromobacter* |
| 1.02E+04 | -1.59E+00 | 6.01E-01 | -2.65E+00 | 8.16E-03 | 8.98E-02 | *Sphingobium* |
| 6.05E+00 | 4.00E+00 | 1.53E+00 | 2.61E+00 | 9.18E-03 | 9.80E-02 | *Flavipsychrobacter* |
| 7.67E+01 | 2.94E+00 | 1.13E+00 | 2.60E+00 | 9.42E-03 | 9.80E-02 | *Archangium* |
| 1.19E+04 | -2.12E+00 | 8.21E-01 | -2.58E+00 | 9.82E-03 | 9.89E-02 | *Enterobacter* |
| 1.29E+01 | -2.64E+00 | 1.03E+00 | -2.58E+00 | 1.00E-02 | 9.89E-02 | *Peribacillus* |
| 5.32E+01 | 2.79E+00 | 1.10E+00 | 2.53E+00 | 1.15E-02 | 1.08E-01 | *Methylovorus* |
| 2.90E+01 | -1.92E+00 | 7.80E-01 | -2.46E+00 | 1.40E-02 | 1.22E-01 | *Candidimonas* |
| 9.40E+01 | -2.37E+00 | 1.01E+00 | -2.35E+00 | 1.89E-02 | 1.58E-01 | *Serratia* |
| 1.54E+01 | -2.60E+00 | 1.12E+00 | -2.32E+00 | 2.04E-02 | 1.64E-01 | *Leclercia* |
| 7.76E+01 | -2.00E+00 | 8.63E-01 | -2.32E+00 | 2.04E-02 | 1.64E-01 | *Rahnella* |
| 8.42E+00 | 3.08E+00 | 1.37E+00 | 2.25E+00 | 2.46E-02 | 1.83E-01 | *Gemmobacter* |
| 4.26E+04 | -2.44E+00 | 1.09E+00 | -2.25E+00 | 2.48E-02 | 1.83E-01 | *Pantoea* |
| 7.68E+02 | -1.68E+00 | 7.58E-01 | -2.22E+00 | 2.63E-02 | 1.89E-01 | *Bordetella* |
| 2.47E+01 | -2.43E+00 | 1.10E+00 | -2.22E+00 | 2.65E-02 | 1.89E-01 | *Tatumella* |
| 5.52E+00 | 3.04E+00 | 1.39E+00 | 2.19E+00 | 2.87E-02 | 1.93E-01 | *Thiomonas* |
| 1.05E+01 | 2.34E+00 | 1.07E+00 | 2.19E+00 | 2.87E-02 | 1.93E-01 | *Gilvimarinus* |
| 3.93E+02 | 1.33E+00 | 6.08E-01 | 2.18E+00 | 2.91E-02 | 1.93E-01 | *Ramlibacter* |
| 3.53E+00 | -5.46E+00 | 2.56E+00 | -2.13E+00 | 3.31E-02 | NA | *Listeria* |
| 1.20E+02 | -1.81E+00 | 8.69E-01 | -2.08E+00 | 3.77E-02 | 2.34E-01 | *Brevundimonas* |
| 3.76E+02 | 1.47E+00 | 7.38E-01 | 1.99E+00 | 4.69E-02 | 2.77E-01 | *Asticcacaulis* |
| 7.80E+02 | 1.58E+00 | 7.95E-01 | 1.99E+00 | 4.71E-02 | 2.77E-01 | *Shinella* |
| 2.22E+01 | 1.84E+00 | 9.27E-01 | 1.98E+00 | 4.76E-02 | 2.77E-01 | *Cytophaga* |
| 9.32E+00 | -1.88E+00 | 9.51E-01 | -1.98E+00 | 4.83E-02 | 2.77E-01 | *Porphyrobacter* |

**Supplementary Table 2: DESeq2 output for the differentially abundant fungal genera in the decaying phyllosphere of *N. benthamiana* plants colonized by *V. dahliae* wild type compared to the *VdAMP3* deletion mutant**

| **Base mean** | **log2FC** | **lfcSE** | **stat** | **pvalue** | **padj** | **Genus** |
| --- | --- | --- | --- | --- | --- | --- |
| 7.20E+01 | -3.14E+00 | 7.70E-01 | -4.08E+00 | 4.50E-05 | 3.87E-03 | *Clavispora* |
| 2.26E+01 | -7.19E+00 | 1.77E+00 | -4.05E+00 | 5.03E-05 | 3.87E-03 | *Cordyceps* |
| 1.55E+01 | -5.85E+00 | 1.51E+00 | -3.86E+00 | 1.11E-04 | 6.00E-03 | *Trichomonascus* |
| 2.64E+01 | -5.10E+00 | 1.34E+00 | -3.80E+00 | 1.43E-04 | 6.00E-03 | *Akanthomyces* |
| 9.44E+03 | -4.13E+00 | 1.15E+00 | -3.60E+00 | 3.19E-04 | 1.02E-02 | *Fusarium* |
| 7.01E+01 | 3.29E+00 | 9.81E-01 | 3.36E+00 | 7.91E-04 | 2.12E-02 | *Terfezia* |
| 2.22E+01 | 3.99E+00 | 1.34E+00 | 2.98E+00 | 2.84E-03 | 4.59E-02 | *Rhodotorula* |
| 1.82E+01 | -2.76E+00 | 9.28E-01 | -2.98E+00 | 2.92E-03 | 4.59E-02 | *Beauveria* |
| 4.87E+01 | -8.30E+00 | 2.86E+00 | -2.90E+00 | 3.71E-03 | 5.29E-02 | *Escovopsis* |
| 1.44E+02 | -2.81E+00 | 9.97E-01 | -2.81E+00 | 4.91E-03 | 6.51E-02 | *Yamadazyma* |
| 1.21E+01 | 3.51E+00 | 1.26E+00 | 2.79E+00 | 5.31E-03 | 6.60E-02 | *Ascosphaera* |
| 5.54E+03 | -3.18E+00 | 1.19E+00 | -2.68E+00 | 7.45E-03 | 8.96E-02 | *Trichoderma* |
| 6.49E+00 | -3.79E+00 | 1.50E+00 | -2.53E+00 | 1.14E-02 | 1.08E-01 | *Xylaria* |
| 5.53E+01 | 2.89E+00 | 1.16E+00 | 2.48E+00 | 1.32E-02 | 1.21E-01 | *Golovinomyces* |
| 1.88E+01 | -2.17E+00 | 8.79E-01 | -2.47E+00 | 1.36E-02 | 1.21E-01 | *Valsa* |
| 5.03E+00 | -3.60E+00 | 1.48E+00 | -2.44E+00 | 1.49E-02 | 1.27E-01 | *Acremonium* |
| 2.18E+00 | -4.77E+00 | 2.04E+00 | -2.34E+00 | 1.94E-02 | NA | *Meyerozyma* |
| 2.92E+00 | -4.17E+00 | 1.79E+00 | -2.33E+00 | 2.00E-02 | NA | *Scedosporium* |
| 4.94E+00 | -3.36E+00 | 1.46E+00 | -2.30E+00 | 2.14E-02 | 1.68E-01 | *Ustilaginoidea* |
| 2.98E+01 | -2.60E+00 | 1.15E+00 | -2.26E+00 | 2.35E-02 | 1.81E-01 | *Debaryomyces* |
| 6.66E+01 | -1.45E+00 | 6.64E-01 | -2.19E+00 | 2.86E-02 | 1.93E-01 | *Phaeoacremonium* |
| 1.58E+01 | 2.43E+00 | 1.15E+00 | 2.11E+00 | 3.46E-02 | 2.26E-01 | *Morchella* |
| 2.10E+01 | -1.97E+00 | 9.38E-01 | -2.11E+00 | 3.52E-02 | 2.26E-01 | *Neonectria* |
| 1.09E+01 | -1.77E+00 | 8.45E-01 | -2.10E+00 | 3.59E-02 | 2.26E-01 | *Spathaspora* |
| 5.98E+00 | -3.07E+00 | 1.51E+00 | -2.04E+00 | 4.12E-02 | 2.52E-01 | *Pochonia* |
| 2.02E+00 | -4.67E+00 | 2.30E+00 | -2.02E+00 | 4.29E-02 | NA | *Millerozyma* |
| 3.54E+00 | -2.82E+00 | 1.41E+00 | -2.00E+00 | 4.52E-02 | NA | *Grosmannia* |
| 3.47E+00 | -2.99E+00 | 1.50E+00 | -1.99E+00 | 4.68E-02 | NA | *Cyberlindnera* |

**Supplementary Table 3: Primers used in this study**

| **Name** | **Sequence (5' --> 3')** | **Application** |
| --- | --- | --- |
| VdAve1_qPCR_Fw | TGTTACCAAAGCAGCACACAAGG | Real-time PCR |
| VdAve1_qPCR_Rv | CCTTATGCCTCGTTCCCTTCCAC | Real-time PCR |
| VdGAPDH_Fw | CGAGTCCACTGGTGTCTTCA | Real-time PCR |
| VdGAPDH_Rv | CCCTCAACGATGGTGAACTT | Real-time PCR |
| VdAMP3_qPCR_Fw | ATGAAGCTCATTTCTGC | Real-time PCR |
| VdAMP3_qPCR_Rv | CTAGTTGCAAATGCACAC | Real-time PCR |
| Chr6g02430_qPCR_Fw | CAGAGCACCACTCACCACAT | Real-time PCR |
| Chr6g02430_qPCR_Rv | ATCAGGAGTGGCGTGAAGTC | Real-time PCR |
| ITS1-Fw | AAAGTTTTAATGGTTCGCTAAGA | Real-time PCR |
| St-Ve1-Rv | CTTGGTCATTTAGAGGAAGTAA | Real-time PCR |
| NbRUB_Fw | TCCGGGTATTAGCAAAAGCGT | Real-time PCR |
| NbRUB_Rv | CCCAAGATCTCGGTCAGAGC | Real-time PCR |
| AtRUB_Fw | GCAAGTGTTGGGTTCAAAGCTGGTG | Real-time PCR |
| AtRUB_Rv | CCAGGTTGAGGAGTTACTCGGAATGCTG | Real-time PCR |
| JR2_VdAMP3_LB_Fw | GGTCTTAAUTTTGAGGGGTTCAGCCGATG | To generate VdAMP3 deletion mutant |
| JR2_VdAMP3_LB_Rv | GGCATTAAUGACGATATGAGTGCTTGCGG | To generate VdAMP3 deletion mutant |
| JR2_VdAMP3_RB_Fw | GGACTTAAUAATGCTTGAGATGACGACGC | To generate VdAMP3 deletion mutant |
| JR2_VdAMP3_RB_Rv | GGGTTTAAUCTGCTCACCAAGCCTCCTTC | To generate VdAMP3 deletion mutant |
| VdAMP3_Comp_Fw | GGGGACAGCTTTCTTGTACAAAGTGGTTTGAGGGGTTCAGCCGATG | To generate VdAMP3 complementation mutant |
| VdAMP3_Comp_Rv | GGGGACAACTTTGTATAATAAAGTTGCTGCTCACCAAGCCTCCTTC | To generate VdAMP3 complementation mutant |
| Promoter_VdAMP3_Fw | CTCGGAATTAACCCTCACTAAAGGGAACAAAAGCTGGAGCTCACA  CAACATCTATGCTTCAGAAGGTGGCAAAAGTG | To generate pVdAMP3::eGFP transformant |
| Promoter_VdAMP3_Rv | ATGATGGCCATGTTATCCTCCTCGCCCTTGCTCACCATATTAATTAA  GATTGATGGTGTCAAGAGGGTCTGGGATATGATTG | To generate pVdAMP3::eGFP transformant |
